## Supplemental Material for "Mechanisms of A-type lamin targeting to nuclear ruptures are disrupted in *LMNA*- and *BANF1*-associated progerias"

| **Primer name** | **Sequence (‘5 to 3’)** |
| --- | --- |
| 1452 | TGGTACGTAGGAATTCGCCACCATGCAGCCTTGGCACG |
| 1453 | ATTCCACAGGGTCGACTTACTTGTACAGCTCGTCCATGCCG |
| 1593 | TGTGGTGGTACGTAGGAATTCGCCACCATGGTGAGCAAG |
| 1651 | CGACTCAGCGGTTTAAACCTACATGATGCTGCAGTT |
| 1833 | TGAAAAACACGATAAAGTTTAAAACATGGTGAGCAAGGGC |
| 1834 | CACACATTCCACAGGGTCGACTTATCTAGATCCGGTGGA |
| 1873 | GCTGTACAAGTCCGGAGAGACCCCGTCCCAG |
| 1874 | CATTCCACAGGGTCGACCTACATGATGCTGCAGTT |
| 1903 | CTTGCTCACCATGTTTAAACTTTATCGTGTTT |
| 1924 | CCAGTGTGGTGGTACGTAGCCACCATGACTACCTCACAGAAGCATCG |
| 1925 | CCGGGCCCTCGAATTCTCAGAGGAAAGCGTCACACCA |
| 1927 | CGCCGGCCGGATCCGCCACCATGACTACCTCACAGAAGCATCGCGATTTCGTGACAGAACCA  ATGGGCGAAAAA |
| 1928 | ATTCCACAGGGTCGACTCAGAGGAAAGC |
| 1998 | GCTTTCCTCTGAGAATTCGCGGGATCAATTCCG |
| 2004 | GTACAAGGACCTCGAGATGACTACCTCACAGAAGCATCG |
| 2005 | ATTCCACAGGGTCGACTCAGAGGAAAGC |
| 2035 | GACTCAGCGGTTTAAACTTACATAATTGCACA |
| 2036 | GACTCAGCGGTTTAAACTCAGCGGCGGCTACCACT |
| 2104 | ATCTCGACATCTCGAGAGCCCTACCTCGCAG |
| 2105 | CGACTCAGCGGTTTAAACCTAGCGTGCGTGCTGTGA |
| 2181 | GCTGTACAAGTCCGGAAGCCCTACCTCGCAG |

Table S1. Primers used to generate plasmids.

Table S2. siRNA oligos used in experiments.

| Target | Species | Catalog Number | Name | Sequence (5’ to 3’) |
| --- | --- | --- | --- | --- |
| NT control |  | D-001810-10-05 | D-001810-01 | UGGUUUACAUGUCGACUAA |
|  |  |  | D-001810-02 | UGGUUUACAUGUUGUGUGA |
|  |  |  | D-001810-03 | UGGUUUACAUGUUUUCUGA |
|  |  |  | D-001810-04 | UGGUUUACAUGUUUUCCUA |
| LaA/C | Mouse | L-040758-00 | J-040758-05 | UUAGGGUGAACUUCGGUGG |
|  |  |  | J-040758-06 | UCAAACUCUCGCUGCUUCC |
|  |  |  | J-040758-07 | UCUUCAUCGACUUCCUCUA |
|  |  |  | J-040758-08 | UUCCUCGCUGUAAAUGUUC |
| BAF | Human | L-011536-02 | J-011536-10 | UACGACAAUAGCAAUCUUU |
|  |  |  | J-011536-11 | UUCAUCUUUCUUUAGCACC |
|  |  |  | J-011536-12 | UGCCACGAAGUCUCGGUGC |
|  |  |  | J-011536-13 | UUGCCCAGGACUUCACCAA |
| LEMD2 | Human | L-017941-02 | J-017941-17 | CUAAAAUAUCGGUGGCGAA |
|  |  |  | J-017941-18 | UCACAGAAGCUGCGACUCU |
|  |  |  | J-017941-19 | CUGAAUUGGUGACGACUGU |
|  |  |  | J-017941-20 | CGAGGAGCGGUUACGGGAA |
| Ankle2 | Human | L-181819-00 | J-181819-12 | AUACUCUCUGAUCCGCUCC |
|  |  |  | J-181819-13 | UUCCCGAGUUGGUCUGCGG |
|  |  |  | J-181819-14 | AACGCUACGUCCUAUAUUU |
|  |  |  | J-181819-15 | AUUCCGAACAACCCAAAUC |
| Emerin | Human | L-011025-00 | J-011025-06 | GUAUCCGAAAGAUCUGCGU |
|  |  |  | J-011025-07 | UACGAGUUGAUCCUACUAC |
|  |  |  | J-011025-08 | UAGGAUAAUAGGACAGGUC |
|  |  |  | J-011025-09 | UGAAACAGGGCGGUAGUGC |
| ZmpSte24 | Human | 430824 | S20066 (sense) | CGAAUAGUUUUGUUUGACAtt |
|  |  |  | (antisense) | UGUCAAACAAAACUAUUCGct |

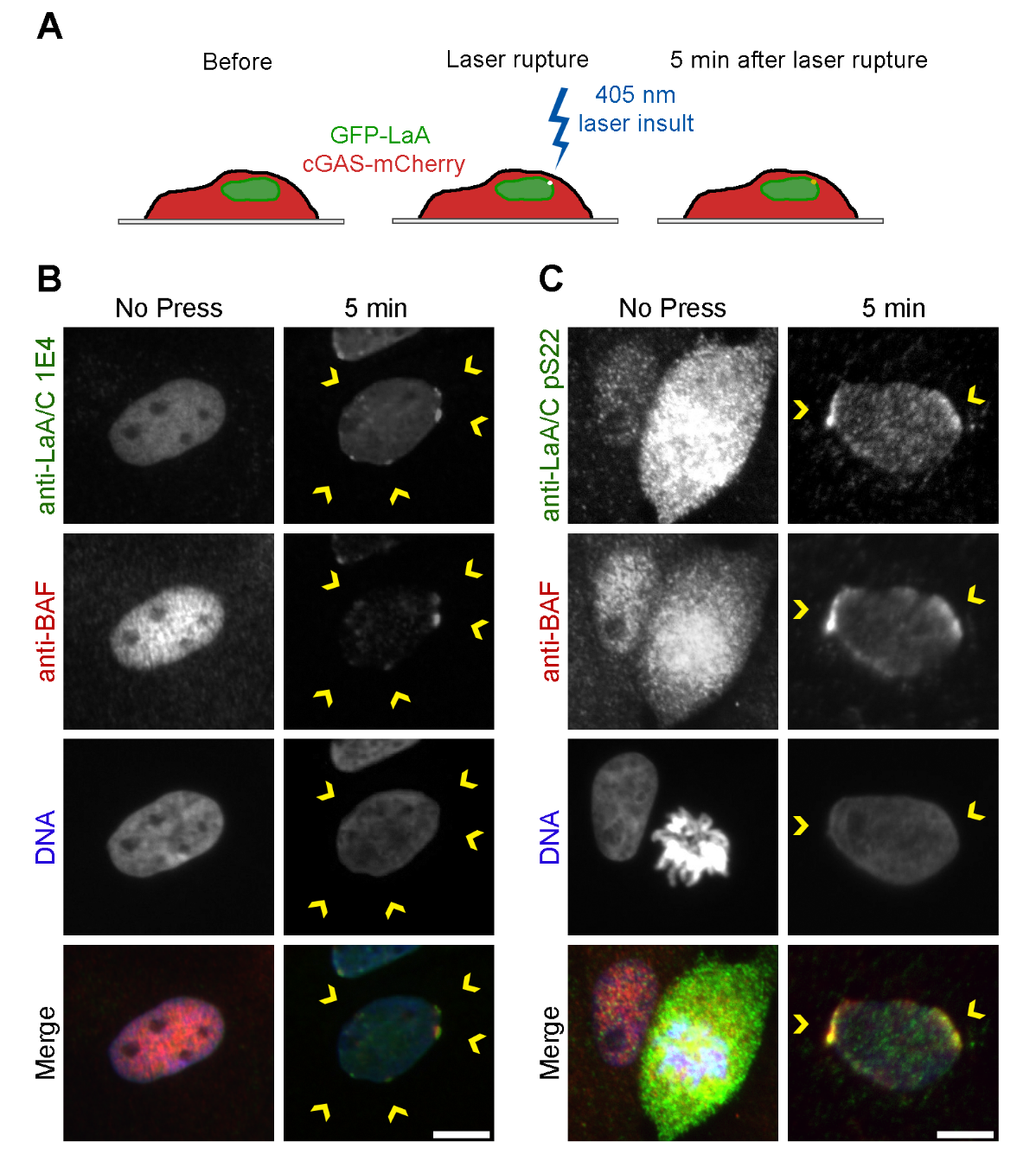

**Supplementary Figure S1**. **Endogenous nucleoplasmic A-type lamins accumulate at nuclear ruptures**. (A) To induce nuclear rupture at a precise location on the nuclear envelope, a 405 laser is applied via a tornado scan to the nuclear envelope for 6-8 sec. MCF10A cells that experienced mechanically-induced nuclear ruptures on via a cell compression chamber were stained to visualize mobile A-type lamins at 5 min post rupture with either (B) the mouse monoclonal antibody 1E4 or (C) an antibody that recognizes A-type lamins phosphorylated at Ser-22. Anti-BAF staining was used as an endogenous rupture reporter. Yellow arrows indicate nuclear rupture locations. Scale bar, 50 μm.

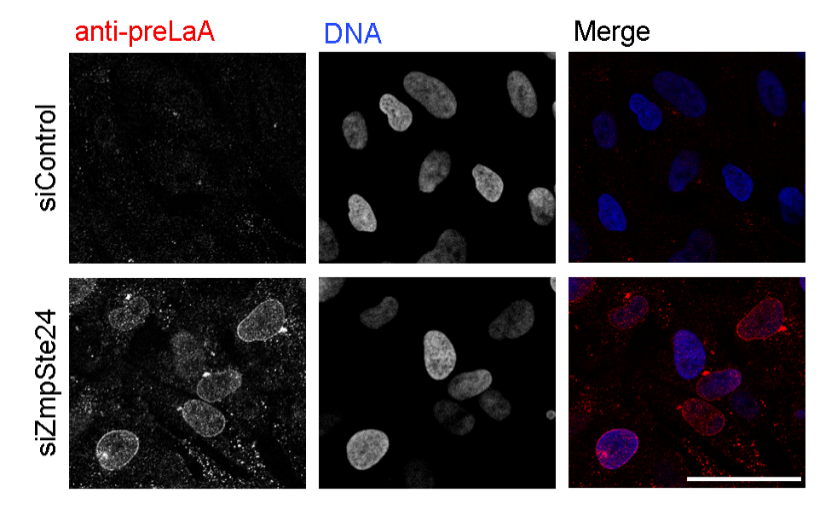

**Supplementary Figure S2. Zmpste24 depletion results in prelamin A accumulation in BJ-5ta cells.** Immunofluorescence confirmation of endogenous prelamin A (preLaA) accumulation in BJ-5ta cells after 144 hr siRNA knockdown of ZmpSte24. Scale bar, 50 μm.

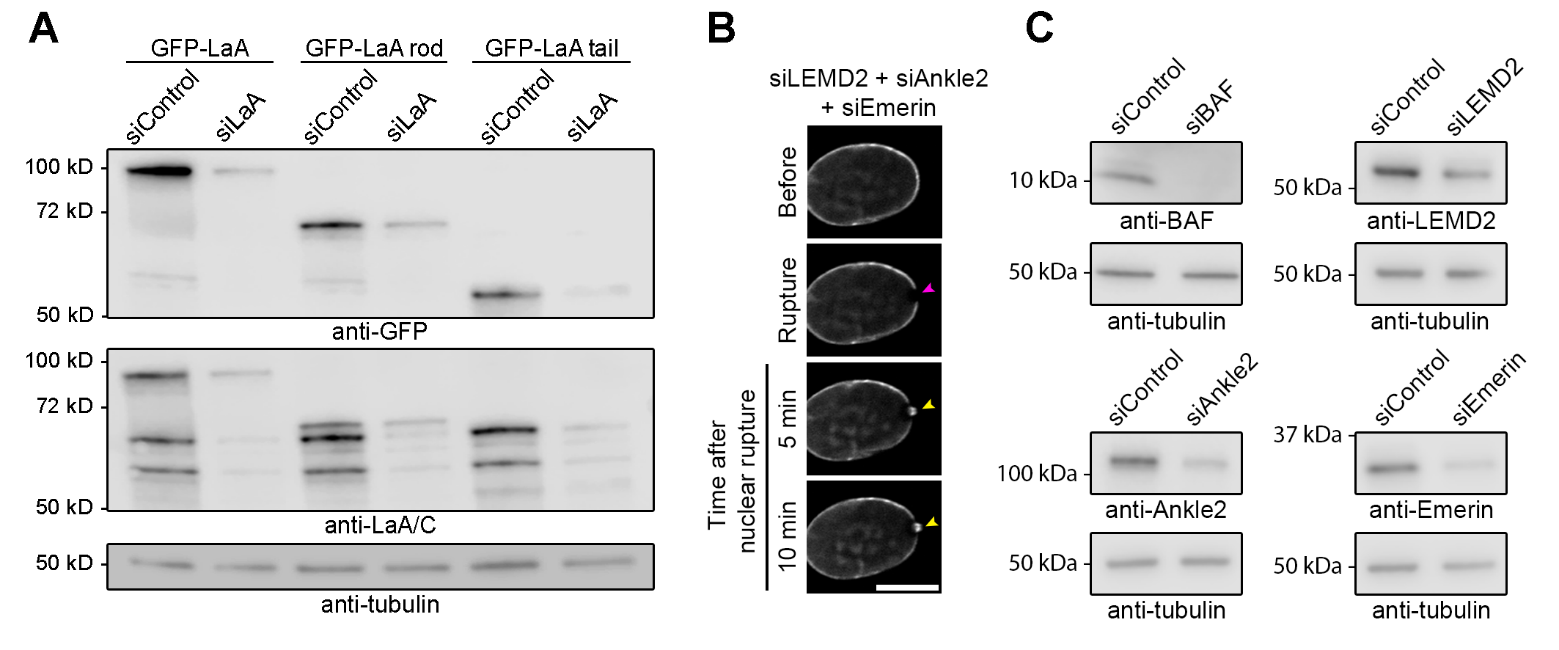

**Supplementary Figure S3. The lamin A tail domain is required for lamin A targeting to nuclear rupture sites.** (A) Representative immunoblot confirmation of endogenous mouse LaA/C depletion after 120 h mouse siLaA/C treatment on NIH3T3 cells stably expressing human GFP-LaA, GFP-LaA rod (aa 1-435), or GFP-LaA tail (aa 391-646). Anti-tubulin was used as a protein loading control. (B) Representative images of laser-induced nuclear rupture events in BJ-5ta cells stably expressing GFP-LaA that underwent a triple siRNA knockdown of the LEM-domain proteins LEMD2, Ankle2, and emerin for 120 h before induction of nuclear rupture. (C) Representative western blot confirmation of 120 h siRNA depletions for BAF, LEMD2, Ankle2, and emerin. Anti-tubulin was used as a protein loading control.

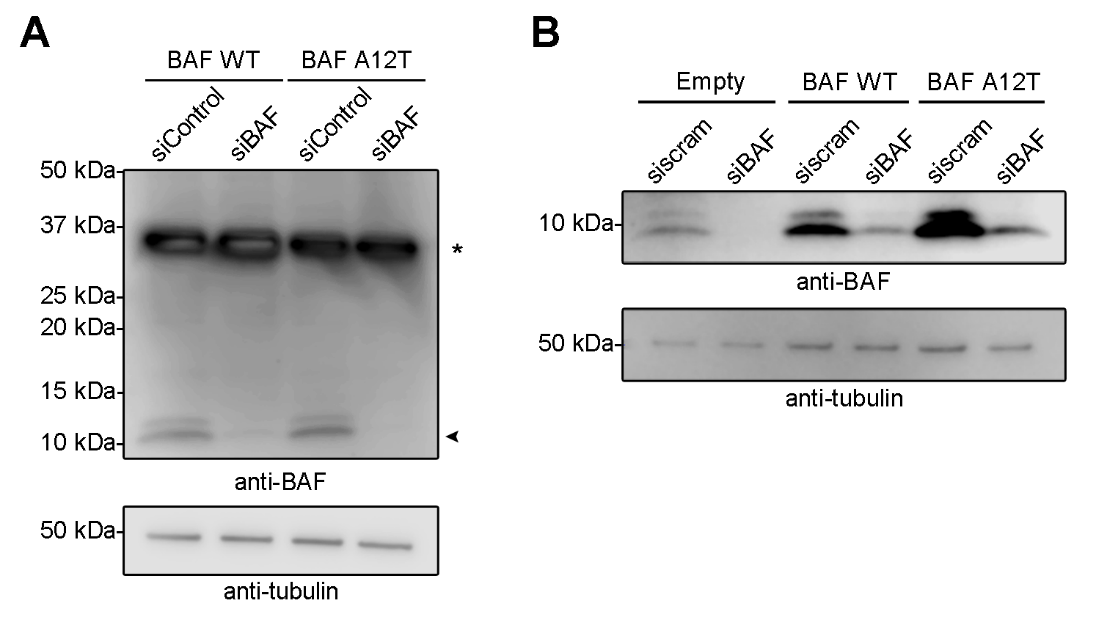

**Supplementary Figure S4. Codon optimized BAF is resistant to siRNA BAF depletion.** (A) Representative immunoblot confirmation of endogenous BAF depletion after 120 h siBAF treatment on BJ-5ta cells stably expressing GFP-tagged codon optimized BAF WT or codon optimized BAF A12T. Asterisk signifies GFP-tagged BAF, and arrowhead signifies endogenous BAF. Anti-tubulin was used as a protein loading control. (B) Representative immunoblot of 120 h siBAF depletion in BJ-5ta cells stably expressing Empty-IRES-GFP-NLS, untagged codon optimized BAF WT-IRES-GFP-NLS, or untagged codon optimized BAF A12T-IRES-GFP-NLS. Anti-tubulin was used as a protein loading control.

**Supplementary Video 1.** Videos of Figure 1B. BJ-5ta cells co-expressing cGAS-mCherry and GFP-LaA (segment 1), GFP-LaC (segment 2), or GFP-LaB1 (segment 3) were monitored for protein accumulation for 10 min after nuclear rupture (yellow arrowhead). Scale bars, 10 μm.

**Supplementary Video 2.** Videos of Figure 2A-B. BJ-5ta cells coexpressing GFP-LaA and cGAS-mCherry underwent siControl (segment 1) or siZmpSte24 (segment 2) treatment for 144 h before laser-induced nuclear rupture and monitored for protein accumulation. (Segment 3) BJ-5ta cells coexpressing GFP-LaA L647R and cGAS-mCherry were monitored for protein accumulation for 10 min after nuclear rupture (yellow arrowhead). Scale bars, 10 μm.

**Supplementary Video 3.** Videos of Figure 2C-D. BJ-5ta cells coexpressing either GFP-LaA L647R (segment 1) or GFP-LaB1 (segment 2) with cGAS-mCherry underwent laser-induced nuclear rupture after 48-72 h incubation with 10 μM farnesyl transferase inhibitor-277 (FTI-277). Protein accumulation was monitored for 10 min after nuclear rupture (yellow arrowhead). Scale bars, 10 μm.

**Supplementary Video 4.** Videos of Figure 3A. NIH3T3 cells stably expressing human GFP-LaA (segment 1), GFP-LaA rod (aa 1-435) (segment 2), or GFP-LaA tail (aa 391-646) (segment 3) and coexpressing cGAS-mCherry were depleted of endogenous mouse LaA/C before undergoing nuclear rupture and monitored for protein accumulation for 10 min following nuclear rupture (yellow arrowhead). Bars, 10 μm.

**Supplementary Video 5.** Videos of Figure 4A. BJ-5ta cells coexpressing GFP-LaA and cGAS-mCherry (segment 1), GFP-LaA Δ50 and cGAS-mCherry (segment 2), or GFP-LaA K542N and cGAS-mCherry (segment 3) were monitored for protein accumulation for 15 min following nuclear rupture (yellow arrowhead). Scale bars, 10 μm.

**Supplementary Video 6**. Videos of Figure 4D. NIH3T3 cells stably expressing human GFP-LaA tail (segment 1), GFP-LaA Δ50 tail (segment 2), or GFP-LaA K542N tail (segment 3) and coexpressing cGAS-mCherry were depleted of endogenous mouse LaA/C before undergoing nuclear rupture and monitored for protein accumulation for 10 min following nuclear rupture (yellow arrowhead). Bars, 10 μm.

**Supplementary Video 7.** Videos of Figure 5A-B. BJ-5ta cells stably expressing GFP-tagged codon optimized BAF WT (segment 1-2) or BAF A12T (segment 3-4) were depleted of endogenous BAF via siRNA transfection before laser-induced nuclear rupture and monitored for protein accumulation for 10 min after nuclear rupture (yellow arrowhead). Bars, 10 μm.

**Supplementary Video 8.** Videos of Figure 5C. BJ-5ta cells stably expressing untagged codon optimized BAF WT-IRES-GFP-NLS and either mCherry-LaA (segment 1-2) or mCherry-LaA tail (segment 3-4) were depleted of endogenous BAF via siRNA transfection before laser-induced nuclear rupture and monitored for protein accumulation for 10 min after nuclear rupture (yellow arrowhead). Bars, 10 μm.

**Supplementary Video 9.** Videos of Figure 5D. BJ-5ta cells stably expressing untagged codon optimized BAF A12T-IRES-GFP-NLS and either mCherry-LaA (segment 1-2) or mCherry-LaA tail (segment 3-4) were depleted of endogenous BAF via siRNA transfection before laser-induced nuclear rupture and monitored for protein accumulation for 10 min after nuclear rupture (yellow arrowhead). Bars, 10 μm.
